# The Vk*MYC Mouse Model recapitulates human multiple myeloma evolution and genomic diversity

**DOI:** 10.1101/2023.07.25.550482

**Authors:** Francesco Maura, David G. Coffey, Caleb K Stein, Esteban Braggio, Bachisio Ziccheddu, Meaghen E Sharik, Megan Du, Yuliza Tofaya Alvarado, Chang-Xin Shi, Yuan Xiao Zhu, Erin W. Meermeier, Gareth J. Morgan, Ola Landgren, P. Leif Bergsagel, Marta Chesi

**Author notes:** **Corresponding authors:** Francesco Maura, University of Miami, 1120 NW 14th Street, Miami, FL 33136, Marta Chesi, Mayo Clinic, 13400 East Shea Boulevard, Scottsdale, AZ 85259. These authors contributed equally (Francesco Maura, David G. Coffey, P. Leif Bergsagel, Marta Chesi).

## Abstract

Despite advancements in profiling multiple myeloma (MM) and its precursor conditions, there is limited information on mechanisms underlying disease progression. Clincal efforts designed to deconvolute such mechanisms are challenged by the long lead time between monoclonal gammopathy and its transformation to MM. MM mouse models represent an opportunity to overcome this temporal limitation. Here, we profile the genomic landscape of 118 genetically engineered Vk*MYC MM and reveal that it recapitulates the genomic heterogenenity and life history of human MM. We observed recurrent copy number alterations, structural variations, chromothripsis, driver mutations, APOBEC mutational activity, and a progressive decrease in immunoglobulin transcription that inversely correlates with proliferation. Moreover, we identified frequent insertional mutagenesis by endogenous retro-elements as a murine specific mechanism to activate NF-kB and IL6 signaling pathways shared with human MM. Despite the increased genomic complexity associated with progression, advanced tumors remain dependent on *MYC* expression, that drives the progression of monoclonal gammopathy to MM.

## INTRODUCTION

Multiple myeloma (MM) is a malignancy of post-germinal center plasma cells (PC) consistently preceded by asymptomatic precursor conditions: monoclonal gammopathy of undetermined significance (MGUS) and smoldering multiple myeloma (SMM).^1–4^ The differentiation between progressive and stable precursor conditions is one of the most important unmet clinical needs in the MM community. The ability to identify individuals at high risk of progression, before clonal expansion and onset of symptoms of end-organ damage, will enable strategies of early prevention.^5^

Leveraging whole genome sequencing (WGS) it has recently been shown that patients with stable precursor conditions are characterized by either absence or lower prevalence of myeloma-defining genomic events, including complex structural variant (SV) events (e.g., chromothripsis), APOBEC mutational activity, mutations in distinct driver genes, and copy-number variation (CNV).^6, 7^ Most of these myeloma-defining genomic events are acquired over time and are often detectable several years before disease progression reflecting a long evolutionary history. Using clock-like single base substitutions (SBS) signatures, SBS1 and SBS5, it was estimated that the temporal development of MM and progressive precursor conditions might be longer than three decades in line with what has been observed across large population studies. ^8^ Such a long lead time makes investigations focusing on the early phase of cancer development a significant challenge.

To overcome this limitation and to investigate the relevant aspects of myelomagenesis, we hypothesized that the clinically faithful, immunocompetent Vk*MYC mouse model would also be relevant to identify the key genomic events of human MM progression.^9^ This genetically engineered model of MM is based on the sporadic AID-induced activation of MYC in a single germinal center (GC) B-cell, in a mouse strain, C57Bl/6, that spontaneously develops monoclonal gammopathy. With age, Vk*MYC mice universally develop a progressive expansion of isotype class-switched, somatically hypermutated, monoclonal PC restricted to the bone marrow (BM), highlighting a dependency on the BM microenvironment for PC growth and survival, as reported for human MM^9^. Clinical myeloma defining end organ damaging events (CRAB: renal impairment, anemia, bone disease) occur only after long latency (usually 70 weeks of age), suggesting that additional mutations, beside MYC dysregulation, are required to induce full malignant transformation^9^. An accelerated disease course is noted when MM tumors harvested from aged, *de novo* Vk*MYC mice are transplanted into syngeneic, non-irradiated C57BL/6 wild type mice.^10^ Despite early evidence from aCGH and a small series of Vk*MYC MM interrogated by scRNA that revealed a complex and heterogenous landscape associated with Vk*MYC MM progression^11, 12^, it is largely unclear if these genomic events recapitulate what is observed in human.

Here, we interrogate the genomic and/or transcriptomic landscape of 128 Vk*MYC MM samples representing 118 unique tumors from 39 *de novo* MM, 63 transplantable lines and 25 tumors capable of growing *in vitro*, to capture the mutations spontaneously selected during disease progression in an unbiased way (**Supplementary Figure 1**). We reveal that phenotypic similarities of the Vk*MYC mouse to human MM are driven by common genomic events, including NFkB activation, APOBEC mutational activity, driver gene mutations, aneuploidies and complex structural variants.

## RESULTS

### Landscape of SNV in driver genes in Vk*MYC MM

Tumor DNA from Vk*MYC MM was analyzed using WGS (n = 41) and WES (n = 27). This cohort included 15 aged mice that spontaneously developed MM (i.e., *de novo*), 33 recipient mice transplanted with tumor cells from *de novo* donors (i.e., transplanted), and 20 *in vitro* cultured tumor cells derived from either *de novo* or transplanted mice (i.e., *in vitro*) (**Supplementary Table 1-3**). Five tumors were interrogated at different stages: four pairs (Vk12598, Vk12653, Vk31159, and Vk37553) were collected from transplanted and *in vitro* samples, and one (Vk36040) from *de novo* and transplanted samples. The overall Vk*MYC MM total mutational burden was 3.8/Mb (range 0.6-20). Importantly, the number of variants per Mb per sample was not significantly different between Vk*MYC WGS and WES indicating a lack of bias in the sequencing technology for detecting coding variants (**Supplementary Figure 2A**).

To identify driver genes under positive selection, the ratio of non-synonymous to synonymous substitutions (dN/dS) was tested using the *dndscv* R ^13^ package, both considering all mutations and restricting to a catalogue of oncodrivers generated by combining known MM driver genes and the COSMIC census.^8, 14, 15^ A total of 11 driver genes under positive selection were detected (q<0.1; **Figure 1A, Supplementary Figure 2B** and **Supplementary Table 4**), six of which are related to known driver genes in MM (*Dusp2, Trp53, Nfkbia, Pim1, Tent5c/Fam46c, H1f4/Hist1h1e*). Positively selected genes not previously described in MM included *H1f2/Hist1h1c*, *Pten*, and *Tbsb4x.* Additional non-synonymous SNVs were found in known human MM driver genes such as *Nras*, *Kras, Cyld*, *Sp140*, *Rasa2,* and *Dis3*, but at a lower frequency compared to human MM (**Figure 1A**). The non-synonymous mutational burden progressively increased across the different mouse stage: *de novo* (median 18 per sample; range 3-39), transplanted (median 43 per sample; range 9-149), and *in vitro* (median 74 per sample; range 19-273) (**Figure 1B**). Interestingly, the nonsynonymous mutational burden of *de novo* samples was similar to human stable precursor conditions and the one observed in transplanted samples was similar to human MM and progressive precursor conditions (**Figure 1B**). *Tp53, Dusp2, and H1f4* mutations were significantly enriched in MM tumors growing *in vitro*, frequently co-occurred in the same sample (p<0.05 using Fisher’s Exact test; **Figure 1C; Supplemental Figure 2C**). This is in line with the almost universal p53 inactivation reported in human MM cell lines.^16, 17^ Additional patterns of co-occurrence between mutated driver genes were observed such as between *Tent5c* and *Hif4* (p<0.05 using Fisher’s exact test; **Supplemental Figure 2C**). Interestingly, the variant allele frequency (VAF) for 10 out of 11 clonal nonsynonymous mutations in *Pten* and seven out of nine in *Trp53* were greater than 75% (**Supplemental Figure 2D**). Although proper characterization of the allele-specific haplotype was not possible since the mouse genome is enriched for homozygous germline SNPs, the fact that only three of these clonal mutations were associated with copy number loss suggests recurrent presence of copy number neutral loss of heterozygosity (CN-LOH). It is notable that the coding and non-coding mutations and deletions of *Pten* are clustered in the first 2kb of the gene, with a preference for T (27/40 SNV involve T), suggesting the involvement of a transcription linked mutational process with preference for T (**Supplementary Figure 2E**).

**Figure 1.**
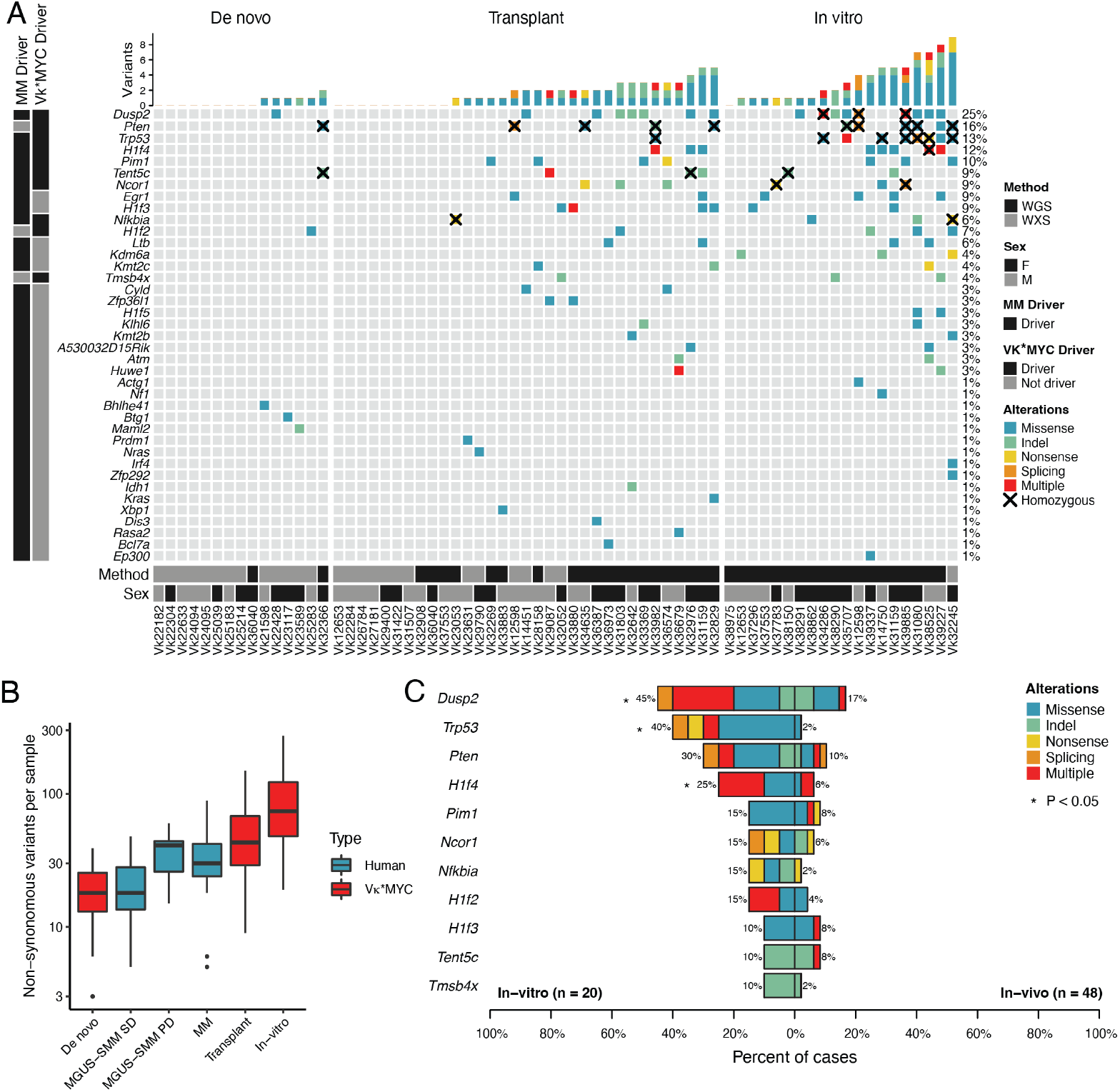
Somatic, non-synonymous single nucleotide variants and indels in driver genes of Vk*MYC MM. **A)** Oncoplot of mutated driver genes detected by the dN/dS method or any nonsynonymous mutations found in a known MM driver genes. **B)** Boxplot comparing non-synonymous mutational burden of Vk*MYC MM and human MGUS, SMM, and MM. **C)** Bar plot showing differentially mutated genes according to Vk*MYC MM type between in vitro and in vivo (i.e., de novo or transplant). P-values were estimated using Fisher exact tests.

### Recurrent copy number alterations in Vk*MYC MM

To characterize the landscape of CNV in Vk*MYC MM, we combined our WES and WGS cohort with additional 28 cases interrogated by aCGH, respectively (**Supplementary Table 5**). For CNV calls in 11 Vk*MYC MM tumors with WES and mate-pair WGS data, we selected the latter. Applying GISTSIC2.0 to 96 Vk*MYC MM we uncovered recurrent arm-level (n=7; 4 gains and 3 deletions) and focal (n=20; 5 gains and 15 deletions excluding the immunoglobulin loci) CNV in the Vk*MYC MM genome (**Supplementary Table 6-7; Figure 2A**). Large chromosomal duplications (i.e., >50% of the chromosome size) and trisomies were observed in 77% of cases with evidence of multiple co-occurring trisomies in 26% cases (**Figure 2B; Supplementary Figure 3A**). To define the temporal relationship between these large chromosomal duplications, we estimated the molecular time of each duplicated segment (i.e., corrected ratio between duplicated and non-duplicated mutations).^14, 18^ Similarly to human hyperdiploid MM, large trisomies in the Vk*MYC myeloma tended to be acquired in the same time window in 8/10 (80%) cases where this analysis was possible (**Figure 2C; Supplementary Table 8)**. However, the Vk*MYC MM molecular time estimates were significantly higher compared to that observed in human MM, suggesting a later acquisition in disease pathogenesis (p<0.00001 using Wilcoxon-test; **Figure 2D**).

**Figure 2.**
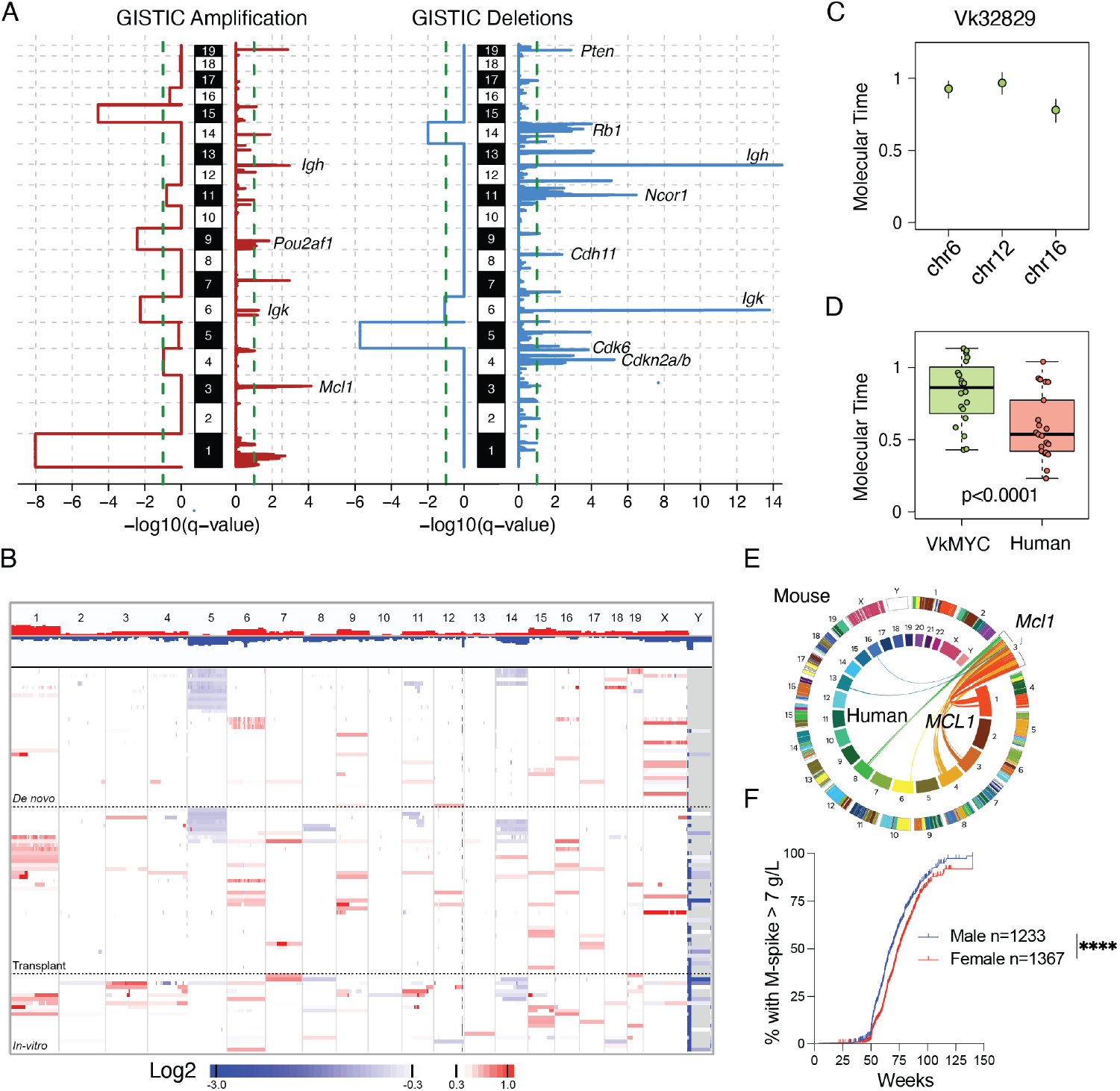
Recurrent focal and broad copy number alterations in Vk*MYC MM. **A)** Significant CNA GISTIC2.0 whole chromosome and focal peaks and involved genes across 96 Vk*MYC MM mice. **B)** Heatmap summarizing the copy number abnormalities of unique Vk*MYC MM, included in the GISTIC analysis. **C)** Example molecular time analysis for the indicated clonal chromosomal gain in one representative Vk*MYC mouse. **D)** Molecular time comparison between Vk*MYC mouse and human MM. P-values were estimated using Wilcoxon text. **E)** Synteny of murine chromosome 3 with human chromosome 1p. **F)** Percentage of male and female Vk*MYC MM with an M-spike greater than 7 g/L, measured every 10 weeks by serum protein electrophoresis. The number of mice analyzed is indicated, as well as the P values calculated by log-rank (Mantel-Cox) test.

As we previously noted, large chromosome 5 loss was observed in 28% of cases confirming the driver role of this large deletion in the Vk*MYC MM.^11, 12^. Moreover, we confirmed chromosome 14 was the second large chromosomal loss that emerged as enriched in our cohort (18.7%). Interestingly, chromosome 14 in mice is largely syntenic with chromosome 13q, the most frequent large chromosomal loss in human MM.^11^ Large and focal gain on chromosome 3 (syngenic with human 1q) and *MCL1* were observed in 12.5% (**Figure 2B,E**). Several known cancer and human MM driver genes were observed within focal GISTIC peaks including homozygous deletions of *Cdkn2a/b* (7.3%), *Rb1* (7.3%), and *Pten* (4%) (**Figure 2A-B, 2E, Supplementary Table 6**). Although 11qB2 (syntenic to human 17p) was detected as recurrently loss by GISTIC, the peak included *Ncor1* but not *Trp53*. As shown above, the lack of significance for *Trp53* (11qB3) CNV loss was due to the fact that all but two *Trp53* clonally mutated cases had a CN-LOH that is not accounted by GISTIC as a loss. Investigating separately CNV on chromosome X, we noted that out of 45 female, 35 (78%) had lost one X chromosome, and out of 44 male, 30 (68%) had lost the Y chromosome. Loss of X and Y chromosomes has been identified by conventional karyotype in human MM, with a frequency of 33% and 20%, respectively^19, 20^. We also confirmed the high prevalence of complete loss of *Kdm6a* (7%), known putative driver gene in human MM, in four male and three female.^11, 21–23^ Interestingly, of those five male, four also lost the *Kdm6a/Utx* homolog *Uty*. Moreover, analysis of M-spike incidence over time identified a highly statistically significant acceleration in tumor development in Vk*MYC male mice compared to female, similar to what has been reported in human MM (**Figure 2F**)^24^.

Overall, our unsupervised CNV analysis identified several similarities between Vk*MYC and human MM, including recurrent events on distinct driver genes and the simultaneous acquisition of large chromosomal duplications and deletions. Further analysis revealed differences in the frequency of genomic abnormalities found across the three stages of Vk*MYC MM progression: *de novo*, transplant and *in vitro* (**Figure 2B**). Focal and large gain (n=21) of chromosome 1 is very rare in *de novo* Vk*MYC MM (5/36, 14%), but is found at higher frequency in the transplant (10/40, 25%) and *in vitro* MM tumors (6/20, 30%). Similarly, chromosome 3 amplification (syntenic to human chromosome 1q) was found in 3 (8.6%) *de novo* tumor, but was detected in 2 (5%) transplants and in 7 (35%) of *in vitro* MM. On the other hand, monosomy 5 is quite common in *de novo* MM (12/36, 33%), but present at lower frequency in transplant (10/40, 25%) and *in vitro* MM (4/20, 20%). Analysis of Vk12653 MM demonstrates that early passages have only one copy of chromosome 5, while late passages and *in vitro* line have two, suggesting a duplication of monosomic chromosome 5 with tumor progression, as recently shown using scRNA data (**Supplementary Figure 3B**).^12^

### Structural variants and complex events

Using WGS (n = 41) and mate-pair WGS (n = 11) we were able to perform the first comprehensive characterization of SV in Vk*MYC MM (**Supplementary Table 1-2 and 9**). After removing the Vk*MYC transgene insertion breakpoints and the deletions associated with VDJ and class switch recombination within immunoglobulin genes (Ig), we observed a median of 8.5 (range 0-351) SVs per tumor, significantly lower compared to what observed in human MM (16, range 0-351; p=0.00017 using Wilcoxon test). Interestingly, evidence of spontaneously acquired chromothripsis, chromoplexy, templated insertion, and complex not otherwise specified SV were observed in 13%, 7.7%, 2% and 23% of mice, respectively (**Figure 3A-D**)^22^.

**Figure 3.**
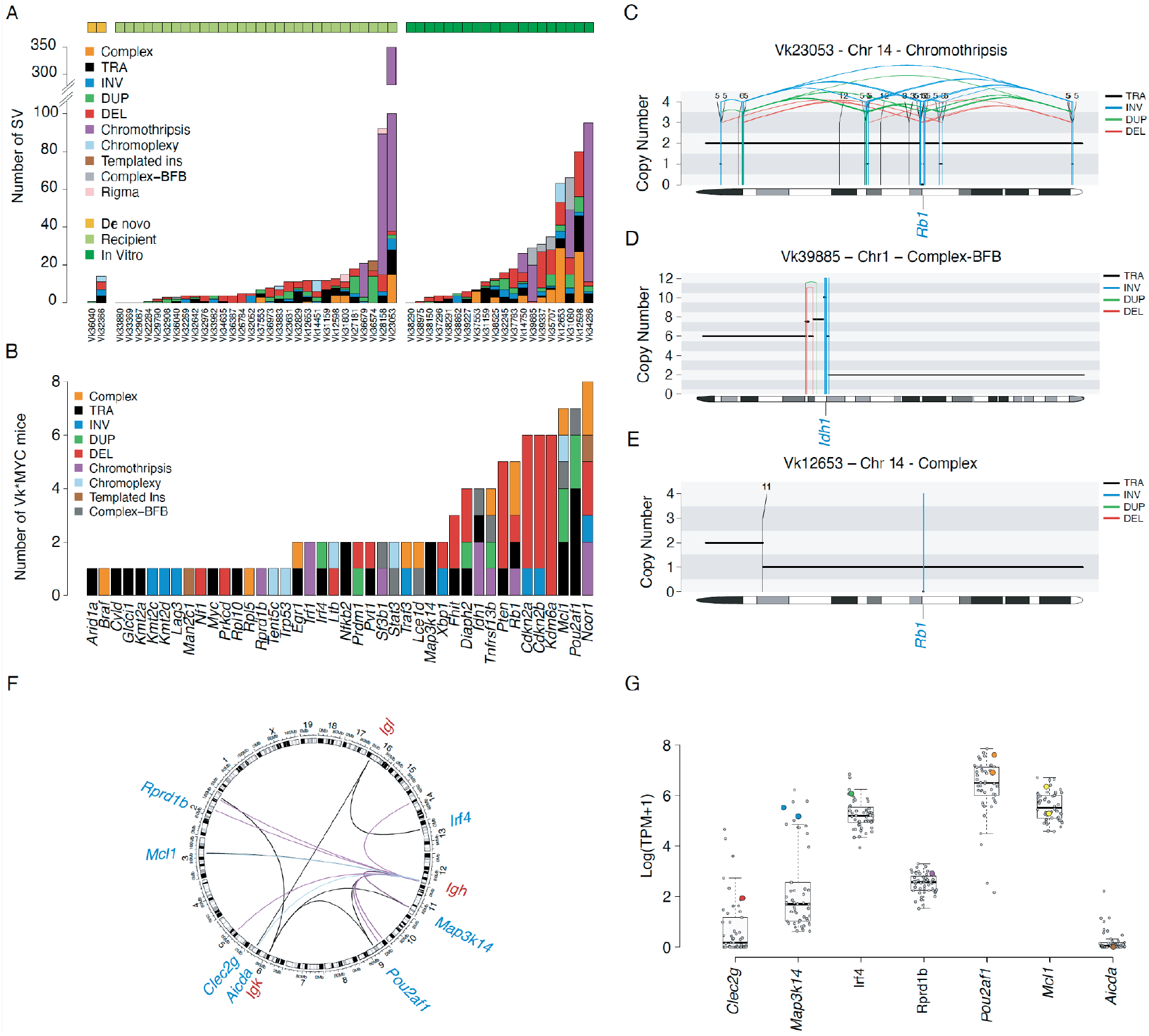
Vk*MYC structural variants (SV) landscape. **A)** Barplot summarizing the prevalence and distribution of SV and complex events across 52 Vk*MYC MM. **B)** Number of Vk*MYC MM mice with SV involving key oncodrivers. **C-E)** Representative examples of SV events involving oncodrivers. BFB: breakage-fusion-bridge. The horizontal black line indicates the total copy number; the dashed orange line indicates the minor copy number. The vertical lines represent SV breakpoints: black: translocations; red: deletions, green: tandem-duplications; blue: inversion. **F)** Circus plot showing all the immunoglobulin translocations detected. **G)** Impact of immunoglobulin translocations on the partners’ gene expression (colored dots).

Complex and single SV events involved several oncodrivers including *Cdkn2a/Cdkn2b*, *Kdm6a*, *Tnfrsf13b*, *Rb1*, *Ncor1* and *Mcl1* (**Figure 3B-E**). Interestingly we observed complex events with evidence of breakage-fusion-bridge inducing multiple focal gains of different oncogenes such as *Mcl1*, *Stat3*, *Pou2af1* and *Idh1* (example shown in **Figure 3D**). As additional similarity between human and Vk*MYC MM, translocations involving the immunoglobulin heavy and light chain loci were detected in eight Vk*MYC MM (15.4%) (**Figure 3F**). Some of these events partnered with known oncogenes promoting their overexpression (e.g. *Map3k14, Irf4*, *Pou2af1,* and *Mcl1*; **Figure 3G**). One translocation creating a fusion gene between *Aicda* and *Igh* was observed, however the expression of the *Aicda* was not detectable by RNA sequencing (RNAseq) in this case (**Figure 3F-G)**.

Overall, we found significant similarities between the SV landscape of human MM and Vk*MYC myeloma, including complex events such as chromothripsis and translocations involving the immunoglobulin genes.

### Retrotransposition of Intracisternal-A-Particle (IAP) LTR mediates gene dysregulation

We have previously reported that retrotransposition of an IAP LTR mediated ectopic expression of *Map3k14* in Vk12653 MM.^25^ Using RNAseq we subsequently detected additional Vk*MYC samples with outlier expression of *Map3k14*, but our pipeline identified a relevant SV involving IgK and IgH in only two of them (**Figure 3F,G**). We therefore visually inspected the BAM files of Vk*MYC MM samples looking for evidence of IAP LTR insertion within the *Map3k14* gene. In 13 tumors *Map3k14* transcription initiated from an ectopic IAP LTR upstream of exon 2 or 3. When available, WGS was able to exactly map the insertion site. (**Figure 4A, Supplementary Table 10**). In all cases with both RNAseq and WGS, outlier expression of Map3k14 could be accounted for by either Ig translocation or IAP insertion. In two cases, lacking WGS, the overexpression of *Map3k14* remained unaccounted. Altogether 17/88 (20%) of the MM tumors analyzed had outlier expression of *Map3k14*. The majority of the LTR insertions result in the elimination of exon 2 encoding the TRAF3 interacting domain known to be important for Map3k14 protein stability, as we and others previously noted in human MM, and cause constitutive NFkB activation, reflected by a high NFkB index by gene expression (**Figure 4B**)^26, 27^. Similarly, high NFkB index was observed in other Vk*MYC MM cases carrying mutations in NFkB regulatory genes described above (i.e., *Nfkbia*, *Ltb*, *Traf3*, *Ikbkb*, *Cyld*) **Figure 4B**).

**Figure 4.**
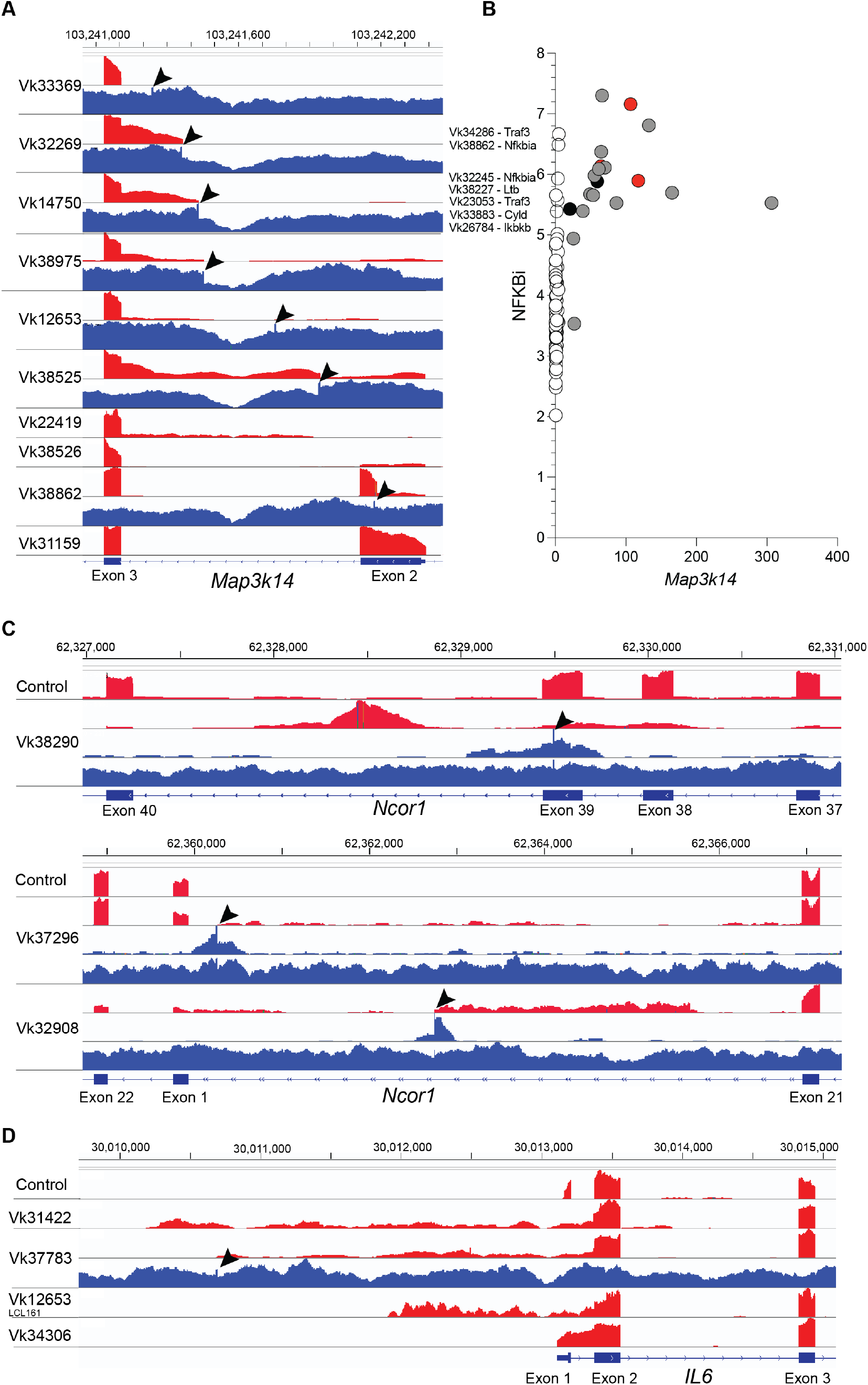
LTR retrotransposition mediated gene dysregulation. **A)** Read depth for RNA (red) and whole genome (blue) sequencing of VkMYC tumor lines for *Map3k14* is shown. By sequence analysis the 5’end of the RNA for these samples originates in an IAP LTR. The IAP insertion site in the DNA is marked by a 6-8 nucleotide amplification indicated by the arrowheads. The RNA for Vk31159 upstream of exon 2 originates from IAP LTR sequences, and WGS identifies an IAP insertion site at chr11:103253336 in intron 1 (not shown). **B)** Gene expression NFkB index plotted versus *Map3k14* expression (RPKM). Each dot represents an individual tumor; in grey are highlighted those with a mapped IAT insertion, in red those with an Ig translocation and in black those with unaccounted *Map3k14* overexpression. The mutated gene in other samples with high NFkB index are listed. **C)** Read depth for RNA (red), discordant reads from whole genome (upper blue), and all reads from whole genome (lower blue) sequencing of VkMYC tumor lines is shown. The IAP insertion site in the DNA is marked by a 6-8 nucleotide amplification indicated by the arrowheads, and is most easily seen visualizing the discordant reads only. Exon numbering for *Ncor1* is from reference transcript NM_011038. The exon labelled 1 is an alternatively spliced exon of *Ncor1* not included in any reference transcripts. It has also been identified as the first exon of *Rip13a/Ncor1*, an *Ncor1* isoform lacking repressor domains. **D)**. Read depth for RNA (red) and whole genome (blue) sequencing of VkMYC tumor lines for *Il6* is shown. By sequence analysis the 5’end of the RNA for these samples originates in an IAP LTR. The IAP insertion site in the DNA is marked by a 6-8 nucleotide amplification indicated by the arrowheads.

We performed an unbiased screen of our WGS using RetroSeq to identify other genes with recurrent insertions of LTR^28^. We identified 38 genes with more than two novel intragenic LTR insertions in the samples analyzed. Of the genes we have identified as recurrently mutated in MM, in addition to *Map3k14*, we found *Ncor1* to have recurrent IAP LTR insertions disrupting its expression, as confirmed by RNAseq (**Figure 4C, Supplementary Table 10**).

Finally, the Balb/c plasmacytoma line MPC11 has been reported to have an IAP LTR insertion upstream of *Il6* resulting in its upregulated expression.^29^ Expression of *Il6* was detectable by RNAseq in only four Vk*MYC MM, which in all cases we determined originated from an IAP LTR (**Figure 4D, Supplementary Table 10).**

### Shared pathways of tumor progression between murine and human MM

Combining the results from the genomic and transcriptomic analyses across all different somatic events spontaneously acquired during Vk*MYC myeloma pathogenesis, we noted the convergence of acquired mutations on pathways activating NFkB (38%), RAS/mTORC1 (27%), cell cycle (48%) and chromatin modifiers (67%) as observed in human MM (**Figure 5A**).^8^ Although we identified classic RAS activating mutations in only two samples (*Nras* Gln61Lys in Vk29790 and *Kras* Gln61Arg in Vk32829), 21 other Vk*MYC tumors (20%) carried *Pten* inactivation, which, like RAS mutations, converges on mTORC1 activation^30^.

**Figure 5.**
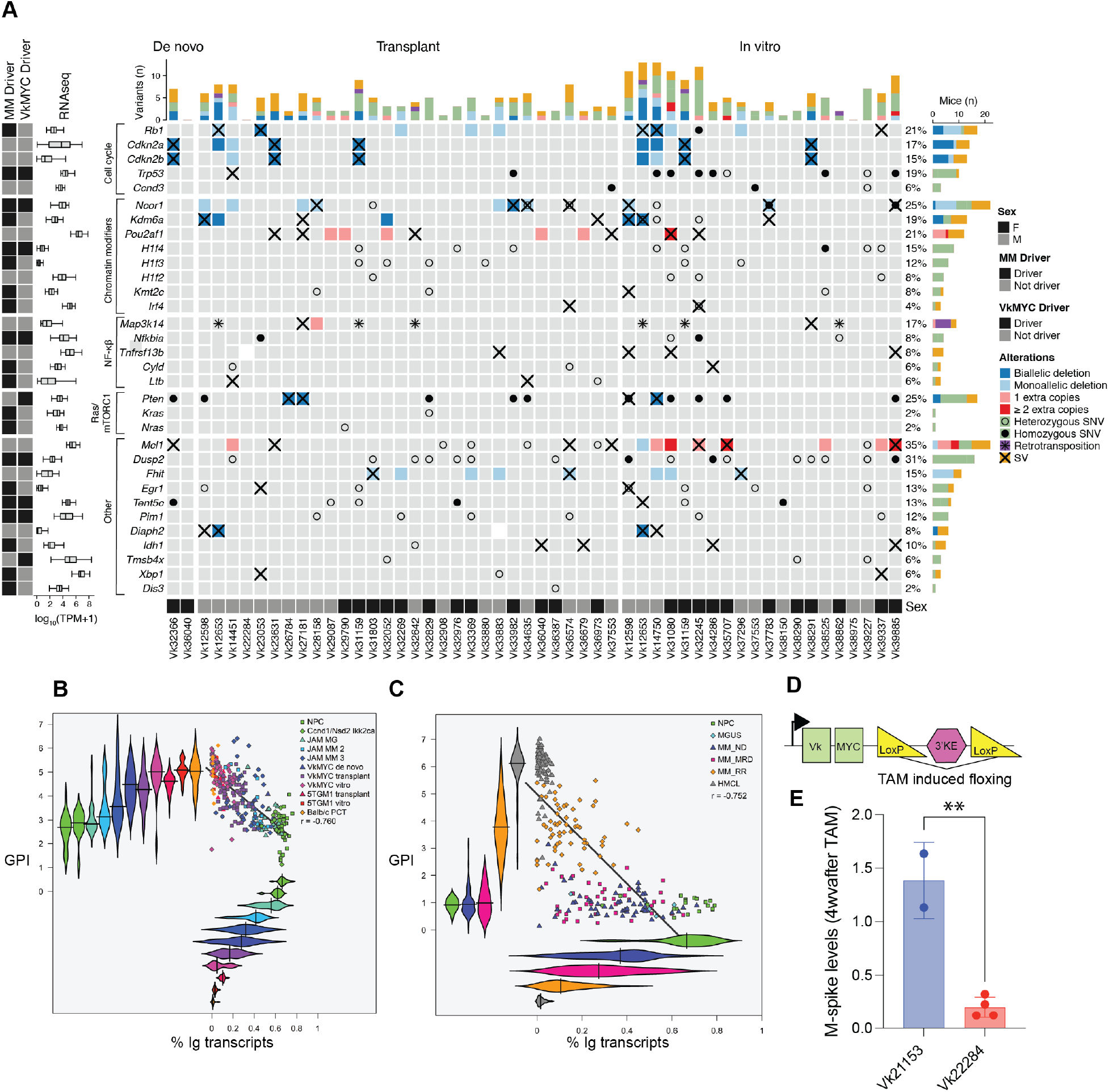
The genomic landscape of the Vk*MYC mouse model of MM. **A)** Heatmap summarizing all the key oncodrivers involved by somatic events in the Vk*MYC MM. Only CNV involving MM oncodrivers and with a length smaller than 3.5 mb are reported. **B-C)** XY scatterplot of percent of immunoglobulin transcription versus gene expression proliferation in murine, **B),** and human, **C),** plasma cell tumors. The % of IG transcription is derived from RNASeq and the GPI score is the mean of the log2 transformed TPMs of the 50-genes comprising GPI^31^. The Pearson correlation score is indicated. **D)** Graphic representation of the Vk*MYC construct (not to scale). Green squares represent the kappa variable region and human MYC exons. The position of the two LoxP sites flanking the 3’ Kappa enhancer is shown, as well as their recombination following tamoxifen induced CRE expression. Horizontal arrow indicates the transcription start point. **E)** M-spike levels measured four weeks after tamoxifen treatment and normalized to day 0 levels in mice bearing Vk21153 Vk*MYCDLox/CreERT2 and Vk22284 Vk*MYC/CreERT2 MM tumors.

As tumors progress from *de novo*, to transplant to *in vitro* stage, we observed an increased genomic complexity which resulted in a higher gene expression proliferation index (GPI), a known adverse event in human MM.^31^ Remarkably, in both human and murine PCs, GPI inversely correlated with immunoglobulin expression: Ig transcription progressively declined from normal PCs, to MGUS – or monoclonal gammopathy decribed in two recent mouse models based on *Ikk2ca activation in germinal center -* ^32, 33^, to newly diagnosed MM – or *de novo* Vk*MYC or the *Ikk2ca* crosses -, to relapse/refractory MM – or transplant Vk*MYC or 5TGM1 MM - to human or murine PC lines that grow *in vitro*, which generally lack Ig expression (**Figure 5B-C**). A tabulation of all the main molecular and behavioral features of each of the 58 established and characterized Vk*MYC transplantable and 25 *vitro* lines is provided (**Supplementary Tables 11-12**)

### Vk*MYC MM tumors remain dependent on MYC expression

Considering the progressive accumulation of genomic abnormalities associated with tumor progression, we wondered if Vk*MYC MM remained dependent on MYC dysregulated expression, which in this model is the driver of progression from monoclonal gammopathy. With this question in mind, we designed the original Vk*MYC transgenic construct with two LoxP site flanking the transgenic 3’ kappa enhancer, enabling its floxing upon CRE mediated recombination, with consequent loss of MYC expression (**Figure 5D**). We subsequently generated a derivative Vk*MYCι1loxP strain, in which the two LoxP sites have been removed, to retain MYC expression in the presence of CRE recombinase. We crossed both Vk*MYC and Vk*MYCι1LoxP mice with a strain carrying a tamoxifen (TAM) inducible CRE allele in the ROSA26 locus (**Supplemental Table 3**). We aged double transgenic mice until they developed MM, and from them established MYC^+^CreERT2^+^ transplantable cell lines: Vk22284 Vk*MYC^+^CreERT2^+^ and Vk21153 Vk*MYC∆LoxP^+^CreERT2^+^ (**Supplementary Figure 4A-C**). Finally, we treated MYC^+^CreERT2^+^ Vk22284 and Vk21153 tumor bearing mice with tamoxifen for five consecutive days, and monitored M-spike levels. While CRE induction had no consequences on Vk21153 MM growth, we observed a rapid reduction of M-spikes in Vk22284 tumor bearing mice, indicating continuous dependency on MYC expression (**Figure 5E**).

### VK*MYC MM SBS signatures landscape

The genome wide mutational landscape of newly diagnosed MM in humans is shaped by 7 different SBS signatures: SBS1 and SBS5 (aging), SBS2 and SBS13 (APOBEC), SBS9 (poly-eta in the germinal center), SBS18 (radical oxygen stress damage), and SBS8. Performing mutational signatures analysis on all VK*MYC MM with available WGS we extracted the same 7 SBS signatures detected in human MM (**Methods; Figure 6A; Supplementary Figure 5A-B; Supplementary Table 13**). Additionally, SBS17, of unknown ethiology, was detected in all but four Vk*MYC MM (n=90%). Presence of SBS9 was also observed in all cases in line with the post-GC origin of the Vk*MYC PC. Looking at the Ig loci and AID-off target genes, we observed clear evidence of somatic hypermutation and SBS84 (AID, a mutational process usually undetectable in mutational signature genome wide analysis) and SBS9 involvement in all but 5 MM (**Figure 6B**), confirmed by a comparative and focused analysis restricted to the clonal Ig genes in murine and human PCs (**Figure 6C**). SBS84 localized mutational activity and AID involvement was also observed across known AID off-target genes, similar to what has been reported in human MM, and implicated in the mutations affecting the driver genes: *Dusp2*, *Pim1*, *H1f4/Hist1h1e, and H1f2/Hist1h1c* (**Supplementary Figure 5**). Consistently, we confirmed targeting of the somatic mutation process to the Vk*MYC transgene causing reversion of the stop codon that controls MYC translation in 71 out of 84 unique MM analyzed by mRNA, associated with an average of 4.5% somatic hypermutation at the immunoglobulin loci ^9^, while the remaining cases without reversion of the stop codon have less than 2% (**Supplementary Tables 11-12**). Moreover, the majority (93.3%) of Vk*MYC MM tumors, like human MM, have undergone class switch recombination to express IgG or IgA; and out of the five cases expressing IgM, four have less than 2% somatic hypermutation at the immunoglobulin loci and lack reversion of the transgenic stop codon, suggesting a germinal center independent tumor origin, similar to the recently reported MM that develops in various different crosses with Ikk2ca activated by IgG1-CRE.^32, 33^ In line with this, the proportion of SBS9 was signficantly lower in IgM VkMYC mice compared to the others (p=0.002 using Wilcoxon test). Interestingly, we noted a shift in the Vk*MYC predominant isotype over time from IgG to IgA, that we suspect is related to a change in the microbiome occurring in our colony (**Figure 6D**).^34^

**Figure 6.**
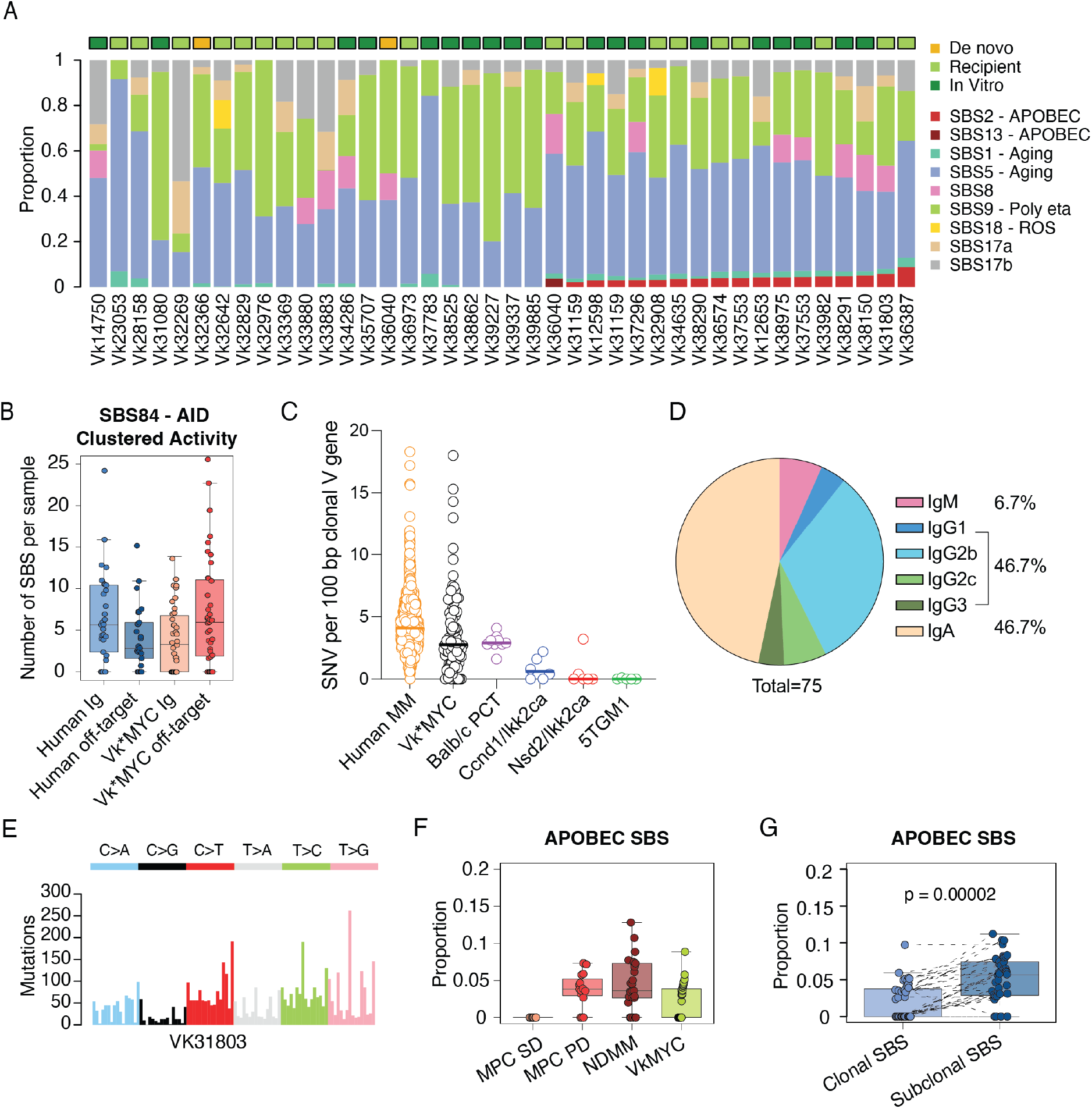
Vk*MYC mouse MM Single base substitutions (SBS) mutational signatures landscape. **A)** Barplot showing the contribution of each mutational signatures for each WGS. **B)** Number of SBS84 (AID) clustered mutations in human and mice involving either immunoglobulin (Ig) or off-target genes. **C)** Percentage of somatic mutation (SNV per 100 base pair) at the productive immunoglobulin allele across tumor types. Each circle represents an individual tumor; in pink are highlighted IgM expressing Vk*MYC MM tumors. **D)** Distribution of immunoglobulin isotypes across Vk*MYC MM tumors. **E)** Example of a 96-classes profile from a Vk*MYC mouse MM with clear APOBEC mutational activity. **F)** APOBEC mutational contribution across Vk*MYC mouse and human MM and precursor conditions. MPC SD: stable myeloma precursor conditions; MPC PD: progressive precursor conditions. **G)** APOBEC mutational activity across clonal and subclonal SBS in Vk*MYC mouse MM. P-values were estimated using paired Wilcoxon text.

Most interestingly, APOBEC mutational activity was detected in 44% of Vk*MYC MM WGS, a proportion slightly lower compared to that observed in human MM and progressive MM precursor conditions (∼80%; **Figure 6E,F**), but in none of the recent *Ikk2ca* based transgenic MM models.^32, 33^ Importantly, no correlation was noted between the presense of APOBEC and AID signatures in line with the distinct nature and role of these two deaminase (**Supplementary Table 12**). To validate the presence of APOBEC mutational signature (SBS2) in Vk*MYC MM we analyzed the SBS signatures on the WES cohort, identifying APOBEC mutational activity in 52% of cases (**Supplementary Figure 6A**). Similar to human MM, APOBEC contribution tends to increase from clonal to subclonal variants, suggesting a late role in Vk*MYC MM development (**Figure 6G**). Additionally, *Apobec3* and *Apobec1* expression were confirmed using RNAseq data from 89 cases (**Supplementary Figure 6B and Supplementary Table 1-2**). No difference in APOBEC mutational contribution was observed between different states, low and high *Apobec3* expression, and *Apobec3* mutated cases. To further investigate the APOBEC role and activity in VK*MYC MM, we interrogated whole bone marrow single cell RNAseq (scRNAseq) data generated from 15 previously published Vk*MYC mice (114,363 total cell count, **Supplementary Figure 7** and **Figure 7**).^12^ *Apobec3* was expressed at single cell level by both, the tumor cells and by normal B-cell and normal plasma cells (**Figure 7E-F**). In contrast *Apobec1* expression was mostly restricted to the tumor plasma cells (**Figure 7C-D**). The fact that immunocompetent Vk*MYC mice is the only model to develop plasma cell tumors with spontaneous aberrant activation of APOBEC further support the similarities in the molecular pathogenesis of disease progression between VK*MYC and human MM.

**Figure 7.**
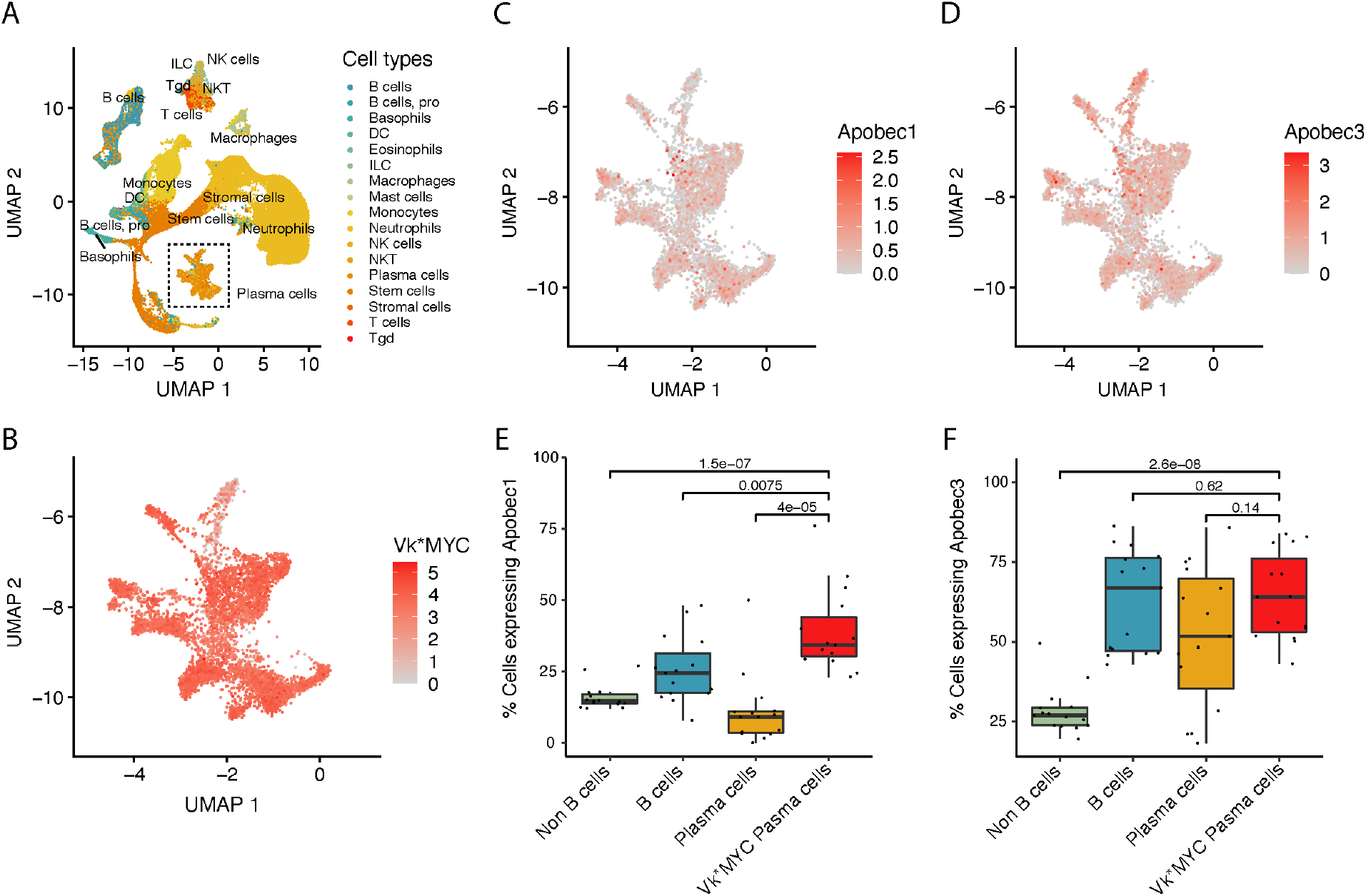
Single cell RNA expression of Apobec1 and Apobec3 across different immune populations in the Vk*MYC mouse. **A)** UMAP analysis of all cells analyzed color coded by cell type. **B)** Expression of Vk*MYC transcript in plasma cells. **C-D)** Expression of Apobec1 (**C**) and Apobec3 (**D**) in plasma cells plasma cells**. E-F)** Difference in Apobec1 (**E**) and Apobec3 (**F**) expression a cross different cell populations. P-values were estimated using Wilcoxon test.

## DISCUSSION

In this study, we performed a comprehensive characterization of the genomic and transcriptomic landscape of Vk*MYC MM, which includes 39 original *de novo* tumors from aged transgenic mice, 63 transplantable tumors lines that can be passaged *in vivo* into non-irradiated, syngeneic C57Bl/6 immunocompetent recipients, and 25 tumors capable of growing *in vitro*. The integration of WGS, WES, bulk and single cell RNA data, revealed the striking genomic similarity between Vk*MYC and human MM. Vk*MYC MM was found to be similar to the 30-40% of human MM characterized by lack of primary IgH translocation and presence *MYC* translocations, also sharing mutations of the NFkB, RAS/mTORC1, cell cycle regulator and chromatin modifier pathways.^14, 15, 26, 35, 36^ These similarities were not just based on individual driver genes or dysregulated pathways but also on broader genomic features, such as presence of chromothripsis, large simultaneously acquired trisomies, and APOBEC mutational activity. Unique to Vk*MYC MM is the hijacking of a murine-specific mutational process that utilizes the IAP transposable element to dysregulate *Map3k14*, *Il6* and *Ncor1*, highlighting the central role of the NFkB, STAT3 and chromatin modifier pathways in MM pathogenesis. IAP expression has been reported to be characteristically high in BALB/c plasmacytomas, with sporadic oncogene activation by retrotransposition.^37^ Here we report three recurrent sites for spontaneous oncogenic insertional mutagenesis by an endogenous retroelement.

Based on our results, we believe the biological fidelity of the Vk*MYC model to human MM is attributable to its unique design. It is based on the introduction of a single *MYC* transgene in the C57BL/6 strain that spontaneously develop monoclonal gammopathy for unknown significance germline predisposition^38^. Remarkably, when backcrossed to a Balb/c strain that does not spontaneously develop gammopathy, Vk*MYC mice fail to develop MM, suggesting that, like in human MM, *MYC* activation is a secondary genomic event that provides the conditions for a transition from a stable to a progressing monoclonal gammopathy^35^. Despite *MYC* activation, Vk*MYC mice remain asymptomatic for at least a year before acquiring MM clinical features, and full MM progression in Vk*MYC mice requires a long latency (70 weeks on average), allowing for the spontaneous acquisition of additional mutations. As a result, each individual Vk*MYC MM displays a unique genomic profile and allows us to identify genomic drivers spontaneously acquired and subsequently selected in an immunocompetent environment. This is in contrast to other genetically modified mouse models of cancer, where engineered expression of multiple oncogenes limits the selection for genomic diversity.^32, 33^ Prior studies identified continued MYC-dependence using mouse models initiatied by MYC.^39^ Here, we show the same holds true in a mouse model where MYC is a progression event. This may in part explain the marked therapeutic efficacy of drugs targeting MYC in the treatment of MM, such as IMiDs.

APOBEC mutational signature is a key genomic feature identified in MM. APOBEC deaminases are known to introduce hundreds of SBS in different cancers, but its aberrant activity is virtually undetectable in normal mouse and human cell genomes.^40^ APOBEC mutational activity has emerged as the most sensitive and prevalent MM defining genomic event associated with MM precursor condition progression.^7, 8, 41^ While *MAF/MAFB* translocated MM patients are characterized by a high APOBEC mutational contribution (i.e., hyper-APOBEC), mostly driven by APOBEC3A isoform, 80% of MM and progressive MM precursor conditions have a lower APOBEC mutational activity where the isoforms 3A and 3B play an equal role (i.e., canonical APOBEC).^8, 42, 43^ Despite its high prevalence in different cancer types ^42^, and recent data suggesting a link with cancer cell immune-escape, genomic instability, and systemic seeding ^44, 45^, the oncogenic role of canonical APOBEC is largely unknown. Moreover, most of the investigations used models in which APOBEC is already constitutively active and mostly driven by the APOBEC3A (hyper-APOBEC),^46, 47^, therefore failing to capture the spontaneous canonical APOBEC activation is seen in MM and lymphomas.^48^ In this context, the ability of the immunocompetent Vk*MYC mouse model to develop a plasma cell tumor with canonical and spontaneous APOBEC mutational activity both represents a confirmation of the genomic similarities between human and Vk*MYC mice MM, as well as a unique model system to explore how and why APOBEC activate during cancer development. Interestingly, leveraging scRNAseq of the entire Vk*MYC plasma cells and surrounding microenvironment, we observed a significant expression of *Apobec3* and a clear enrichment of *Apobec1* compared to other normal population. While *Apobec3* mutational activity has been demonstrated in mice, *Apobec1* emerged as tumor plasma cell specific potential promoter of SBS2 and SBS13.

Interestingly, the genomic life-history of the Vk*MYC MM pathogenesis shared similar patterns with what is observed in human MM. Specifically, MM in both species is characterized by progressively increased proliferation and decrease in immunoglobulin expression, APOBEC mutational activity, mutations in distinct driver genes (e.g. *TP53*), and by the intermediate acquisition of chromothripsis, structural variants and focal copy numbers.^14, 22, 49^ In contrast, certain events co-occurred in both species, but with an opposite timing. In human MM the simultaneous acquisition of large chromosomal gains is usually one of the earliest events, and *MYC* translocation occurs as one of the latest events driving the final progression into symptomatic MM. In contrast to this timeline, *MYC* activation is the earliest events in Vk*MYC MM, and the simultaneous acquisition of large chromosomal gains is usually acquired later in time.^8, 14, 36, 50, 51^ Despite this opposite timeline, our data suggest that the overexpression of *MYC*, alteration of the NFkB pathway and large chromosomal gains are essential for the development of a large fraction of PC tumors in both the Vk*MYC mouse and in humans.

A striking difference between Vk*MYC and human MM is the high prevalence of *Pten* inactivation and low incidence of RAS activating mutation seen in the mouse, that is the exact opposite of what found in human MM. Since both mutations converge on MTORC1, we believe our finding highlights the importance of the MTORC pathway and suggests that the primary importance of RAS mutations in MM may be more through this pathway than MAPK, supported in part by the disappointing effect of MAPK inhibitors in the treatment of RAS activated MM^52^.

Vk*MYC MM cell lines Vk12598 and Vk12653 have been extensively used for myeloma research. Some of the advantages they offer are a high level of immunoglobulin transcription, dependence on the *in vivo* mivroenvironment, rapid engraftment in immunocompetent C57Bl6 mice and fidelity to human MM. The additional cell lines we are reporting here, with their extensive genomic characterization, greatly expands the repertoire of unique MM tumor models that may be used to reflect different aspects of MM biology and we hope will be a rich resource for investigators in the field for years to come.

In summary, through comprehensively genomic profiling of plasma cell DNA and bulk and single cell RNA, the Vk*MYC mouse emerges as a model that recapitulate the genomic profile of a large group of MM. As a result, the Vk*MYC model provides a unique opportunity to uncover the step-wise genomic events within plasma cells not otherwise feasible in human studies to be able define the genomic underpinnings of MM disease progression. Finally, this study highlights the importance and advantage of applying comprehensive genomic investigations in mouse models in which genomic events are spontaneously acquired over time. Detailed knowledge of the spontaneous cancer evolution in such models has major potential in deciphering the early phase of cancer initiations, the selection/acquisition of distinct genomic events in the context of an immunocompetent environment, creating robust biological rational for effective pre-clinical interventions.

## METHODS

### Vk*MYC mouse model and cell lines

All experiments were performed under the approval of the Mayo Foundation Institutional Animal Care and Use Committee and conformed to all the regulatory environmental safety standards. The generation and initial characterization of the Vk*MYC (RRID:MMRC_68098_MU), their derivative lacking LoxP sites, Vk*MYCDLox (RRID: MMRRC_068099-MU) and Vk*MYC^hCRBN^ mice and derived transplantable lines have been reported previously ^9, 53^. *De novo* mice were aged and monitored for tumor burden by serum protein electrophoresis as previously described^11^. M-spikes were quantified by calculating the ratio of densitometric values of the M-spike and albumin bands using the albumin reference value of 27 g/L^54^. Transplantable Vk*MYC lines mice were generated by serial transplantation (at least three times) of approximately 1 million unsorted tumor cells harvested from Vk*MYC *de novo* mice into 4–12 week old C57BL/6 recipient mice of both sexes^53^. Vk*MYC *vitro* cell lines were generated by culturing permissive Vk*MYC cells harvested at the time on necropsy in RPMI-1640 + 10% fetal bovine serum supplemented with glutamine, penicillin, and streptomycin. All cell lines were tested for mycoplasma contamination biannually using the MycoAlert kit (Promega) and were periodically validated by copy-number polymorphism by PCR. Floxing of the transgenic 3’ kappa enhancer was achieved by treating tumor bearing mice with 1 mg tamoxifen (Sigma-Aldrich) in corn oil given by daily i.p. injection for five consecutive days.

### Whole genome, whole exome and mate pair sequencing of VΚ*MYC tumor cells

Myeloma cells were harvested and purified as previously described ^53^. DNA was extracted from isolated plasma cells using Puregene (Qiagen). For WES, paired-end libraries are prepared following the manufacturer’s protocol (Agilent) using the Bravo liquid handler from Agilent. Briefly, 1 ug of genomic DNA was fragmented to 150-200 bp using the Covaris E210 sonicator. The ends were repaired and an “A” base was added to the 3’ ends. Paired end Index DNA adaptors (Agilent) with a single “T” base overhang at the 3’ end was ligated and the resulting constructs were purified using AMPure SPRI beads (Agencourt). The adapter-modified DNA fragments were enriched by 4 cycles of PCR using SureSelect forward and SureSelect ILM Pre-Capture Indexing reverse (Agilent) primers. The concentration and size distribution of the libraries is determined on an Agilent Bioanalyzer DNA 1000 chip.

Whole exome capture was carried out using the Agilent SureSelect Mouse all exon kit. 750 ng of the prepped library was incubated with whole exon biotinylated RNA capture baits supplied in the kit for 24 hours at 65 °C. The captured DNA:RNA hybrids were recovered using Dynabeads MyOne Streptavidin T1 (Dynal). The DNA was eluted from the beads and desalted using purified using Ampure XP beads (Agencourt). The purified capture products was then amplified using the SureSelect Post-Capture Indexing forward and Index PCR reverse primers (Agilent) for 12 cycles. The concentration and size distribution of the libraries was determined on Qubit (Invitrogen) and Agilent Bioanalyzer DNA 1000 chip. Exome libraries were loaded one sample per lane onto Illumina TruSeq v3 paired end flow cells at concentrations of 9 pM to generate cluster densities of 600,000-800,000/mm^2^ following Illumina’s standard protocol using the Illumina cBot and TruSeq Rapid Paired end cluster kit version 3. Some whole-exome libraries were sequenced on a NovoseqPE150 sequencer (Illumina) to generate 12G of raw data per sample. Whole-genome libraries were prepared using the NEBNext® Ultra™ DNA Library Prep Kit (New England Biolabs) and sequenced on a NovoseqPE150 sequencer (Illumina) to generate 75G of raw data per sample.

Nextera Mate Pair libraries were prepared following the manufacturer’s protocol (Illumina). 1ug of genomic DNA in 76 ul EB buffer was simultaneously fragmented and tagged with a biotinylated mate pair junction adaptor. The resulting construct contained a short single stranded sequence gap which was repaired enzymatically according to manufacturer’s protocol (Illumina).The repaired DNA was purified and smaller DNA fragments (< 1500 bp) were removed using AMPure Beads. The size selected fragments were circularization by blunt end ligation for 16 hrs at 30°C using circularization ligase (Illumina). Non-circularized fragments were eliminated by DNA exonuclease treatment. The remaining circularized DNA was again fragmented, this time using the Covaris E210, generating double-stranded DNA fragments with fragment sizes in the 200-2000 bp range. The biotinylated DNA fragments were purified using Dynalbeads M-280 streptavidin beads (Invitrogen) as outlined in the Illumina Mate-Pair protocol. Illumina indexed adapters were added to the DNA on the M-280 beads using the TruSeq Library Sample Preparation kit (Illumina) as follows. The ends of the biotinylated fragments immobilized on the beads were repaired and phosphorylated using Klenow, T4 DNA polymerase, and T4 polynucleotide kinase; after which an “A” base was added to the 3’ ends of double-stranded DNA using Klenow exo- (3’ to 5’ exo minus). Paired end DNA adaptors (Illumina) with a single “T” base overhang at the 3’ end were ligated and the immobilized adapter-modified DNA fragments are enriched by 10 cycles of PCR. The PCR supernatant was recovered from the beads using a magnetic rack. The PCR enriched constructs were cleaned up with AMPure xp beads recovering DNA fragments of approximately 300 - 2000 bp. Concentration and size distribution of the libraries were determined on an Agilent Bioanalyzer DNA 1000 chip and Qubit dsDNA assay (Invitrogen).Libraries were sequenced at 4 samples/lane to generate ∼ 150 million reads/sample following Illumina’s standard protocol using the Illumina cBot 3000/4000 PE Cluster Kit. The flow cells were sequenced as 150 × 2 paired end reads on an Illumina HiSeq 4000 using TruSeq SBS sequencing kit version 3 and HCS v3.3.20 data collection software. Base-calling was performed using Illumina’s RTA version 2.5.2.

Whole genomic and whole exome sequencing reads were aligned the mouse reference genome (GRCm38/mm10) using BWA (Supplemental Figure 10). Base quality score recalibration was performed using GATK tools and duplicates were marked using Picard. Somatic variants were called using Mutect2 and annotated using Annovar. Applying additional post-processing filters, only mutations with a ROQ score (i.e., phred-scaled qualities that alt allele are not due to read orientation artifact) > 89, TLOD score (i.e., log 10 likelihood ratio score of variant existing versus not existing) > 6, and DP (approximate read depth) score > 5 were included in our analysis. Variants in known MM driver genes not passing these quality filters were manual inspected in IGV and were kept within the analysis if they did not appear to be an artifact. Additionaly, we eliminated any WGS samples if there was less than 500 variants per genome or WES samples if there were less than 10 variants per exome.

CNV were called using the GATK4 Somatic CNV pipeline modified for mouse. GISTIC2.0 was used to estimate CNV enriched more than what it would be expect by chance. Focal CNV segments < 10 kb with identical start or end positions in > 3 mice were manually inspected and annotated as polymorphism artifacts and excluded from the analysis. All SV calls are available in **Supplementary Table 5**.

SV were called using Lumpy ^55^ as implemented in Smoove release 19 (https://github.com/brentp/smoove) Complex SV were annotated by manual inspection following the previously published criteria^22^. SV gene involvement is defined by: the genes is within deletions and duplications <3 Mb, the genes is 100Kb from any SV breakpoint, the gene is up to 500Kb from a translocation or inversion. All SV calls are available in **Supplementary Table 11**.

Mutational signatures were estimated using first sigprofiler and *hdp* as de novo extraction, and then mmsig as fitting ^56, 57^. Molecular time analysis was run as previously described (https://github.com/UM-Myeloma-Genomics/mol_time).8

Datasets are available on BioProject PRJNA938752

### aCGH of VΚ*MYC tumor cells

High-resolution aCGH was performed on Gentra-Puregene-cell-kit (Qiagen) purified DNA from 27 mice with the Sureprint G3 mouse CGH 244K microarray kit (Agilent Technologies), as previously described^11^. The data have been submitted with GEO submission #GSE110954.

### Gene expression profiling

RNA from CD138-selected Vk*MYC plasma cells was extracted from TRIzol and further purified on Purelink Micro-Mini Columns (Invitrogen), with an On-Column DNAse Digestion Step. Gene expression profiling was performed on the Affymetrix mouse 430 2.0 array following the manufacturer’s suggested protocol, as previously described^11^. Datasets have been deposited to GEO under submission #GSE111921. For RNAseq, mRNA libraries were prepared using NEBNext Ultra II RNA Library Prep Kit (New England Biolabs) with polyA enrichment and sequenced on a Novoseq S4 sequencer (Illumina) to generate 6G of raw data per sample. RNA sequencing reads were aligned to the mouse reference genome (CRCm38/mm10) using STAR and transcript per million (TPM) gene expression values were obtained using Salmon. Datasets are available Datasets are available on BioProject PRJNA938752. For comparisons between human and murine gene expression, CoMMpass IA19 release was used--limited to bone marrow samples with available RNA-Seq data and either a ‘baseline’ or ‘confirm progression’ reason for visit. The unstranded, salmon TPM counts with immunoglobulin filtering were log2 scaled and used to derive the 50-gene published proliferation score (GPI) and 11-gene published NFkB Index (NFkBi)^27, 31^. Details on immunoglobulin involvement were leveraged from RNA-Seq QC metrics. For mouse comparisons, immunoglobulin filtered RNA-Seq TPM gene counts were similarly used. The GPI and NFkBi gene scores were computed from their respective mouse analogue genes.

### Single-cell RNA sequencing of Vk*MYC bone marrow

scRNAseq data from a previously published study of unsorted Vk*MYC bone marrow mononuclear cells were downloaded from the Sequence Read Archive (SRP214856)^12^. BAM files were converted to fastq format and realigned to a custom version of the mm10 mouse reference genome (refdata-gex-mm10-2020-A) incorporating the Vk*MYC transgene using Cellranger v7.0 (10X Genomics) ^12^. Gene count matrices and sample aggregation was performed using Cellranger. Cells were classified into cell type by SingleR v1.8.1 using the “immgen” reference dataset (**Supplementary Figure 8**). Cell counts were normalized, variable features were found, and dimensionality reduction using PCA and UMAP were performed using Seurat v4.1.1. Plasma cells were identified by as those B cells classified by SingleR and having an average expression score > 0 of selected plasma cell genes (*Sdc1, Tnfrsf17, Slamf7, Xbp1*) using the Seurat AddModuleScore function. Tumor plasma cells were identified as those cells classified as plasma cells and having a Vk*MYC transgene average expression score > 0.

## AUTHOR CONTRIBUTIONS

- Conceptualization: FM, MC, LB
- Data curation: FM, MC, LB, DC
- Formal Analysis: FM, MC, LB, DC
- Funding acquisition: FM, MC, LB
- Investigation: FM, MC, LB, DC, EB, CKS, BZ, MES, MD, EWM, GJM, OL
- Writing – original draft: FM, MC, LB, DC
- Writing – review & editing: all authors

## ACKNOWLEDGMENTS

This work was supported by the National Institute of Health (RO1CA234181, U54CA224018 and CA186781 MC, PLB, CKS, MES, MD, EWM), the Multiple Myeloma Research Foundation (MMRF), the Perelman Family Foundation, the Riney Family Multiple Myeloma Research Program Fund, and by a Sylvester Comprehensive Cancer Center NCI Core Grant (P30 CA 240139).

FM is supported by the American Society of Hematology.

GJM received grant support through a Translational Research Program award from the Leukemia & Lymphoma Society (6020-20).

## CONFLICT OF INTEREST

M.C and P.L.B receive royalties from Vk*MYC, Vk*MYC^hCRBN^ mice and derivative transplantable lines.

## Notes

### Competing Interest Statement

The authors have declared no competing interest.

