## Supplemental Figures for "The Vk*MYC Mouse Model recapitulates human multiple myeloma evolution and genomic diversity"

### SUPPLMENTAL FIGURES

### Vκ*MYC mouse model recapitulates the key myeloma defining genomic events and the life history of multiple myeloma.

**Supplemental Figure 1. Overview of methods.** Tumor DNA was analyzed from 96 Vκ*MYC mice which included 36 mice that developed clonal plasma cell expansion de novo, 40 recipient mice transplanted with tumor cells from *de novo* donors, and 20 *in vitro* cultured tumor cells derived from transplanted mice (**Figure 1, Supplemental Table 1**). DNA was analyzed by whole genome sequencing (WGS), whole exome sequencing (WES), mate pate whole genome sequencing, and array comparative genomic hybridization (aCGH) for single nucleotide variants (SNVs), copy number alterations (CNAs), and structural variants (SV). Tumor cell gene expression (GE) was analyzed by RNA sequencing in 55 Vκ*MYC MM.

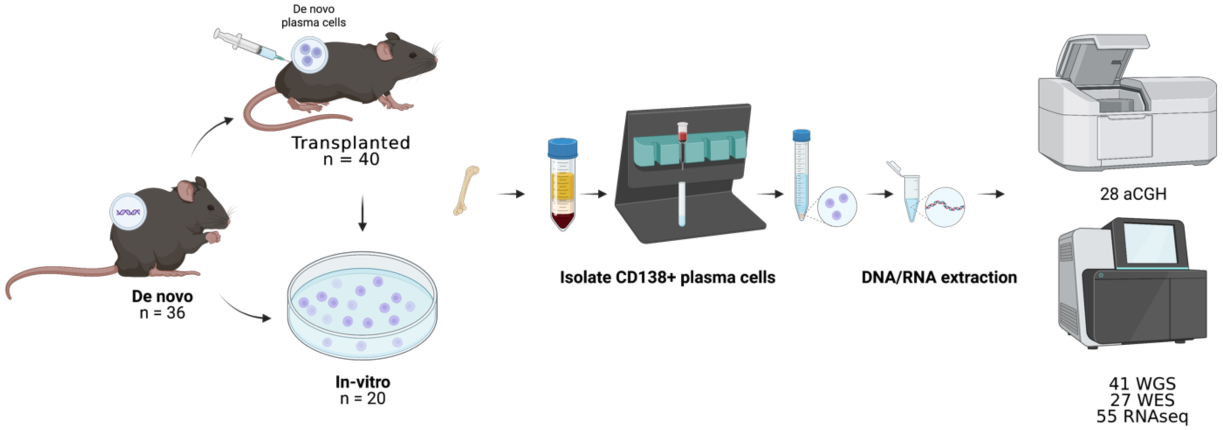

**Supplemental Figure 2. Vk*MYC multiple myeloma (MM) mutational landscape. A)** Similar mutational burden between WGS and WES Vk*MYC MM. **B)** Driver genes extracted by *dndscv* with significant q-value (i.e. <0.01). **C)** Patterns of co-occurrence and mutually exclusivity between mutations in driver genes. **D)** VAF of all nonsynonymous driver mutations**. E)** Distribution of non-coding and coding mutations across *Pten* promoter region.

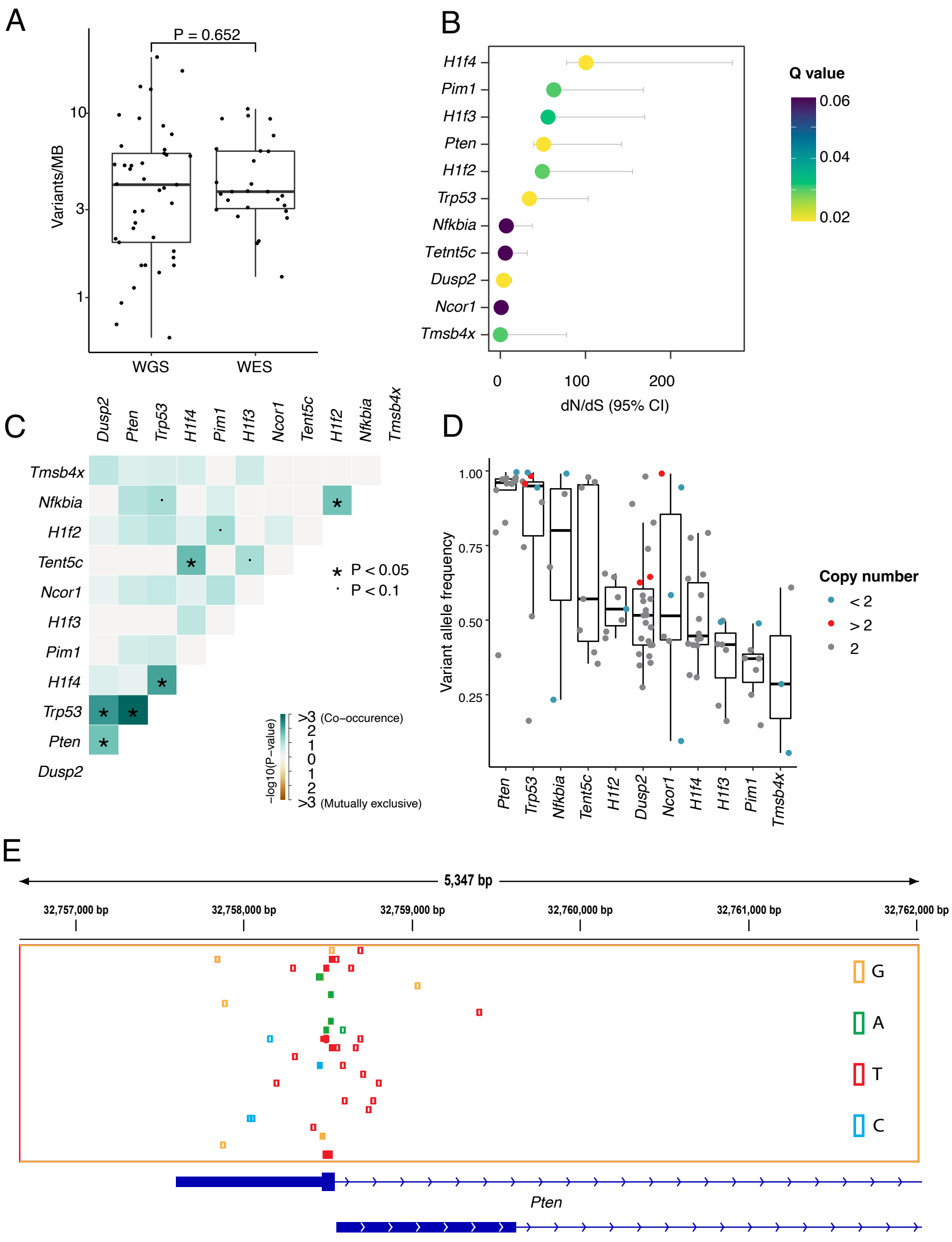

**Supplementary Figure 3. A)** Recurrent copy number alterations involving focal GISTIC peaks in Vk*MYC MM **B)** IGV representation of copy number changes detected over time in Vk12653 MM before and after treatment with bortezomib (BORT) (Chesi et al Blood 2012).

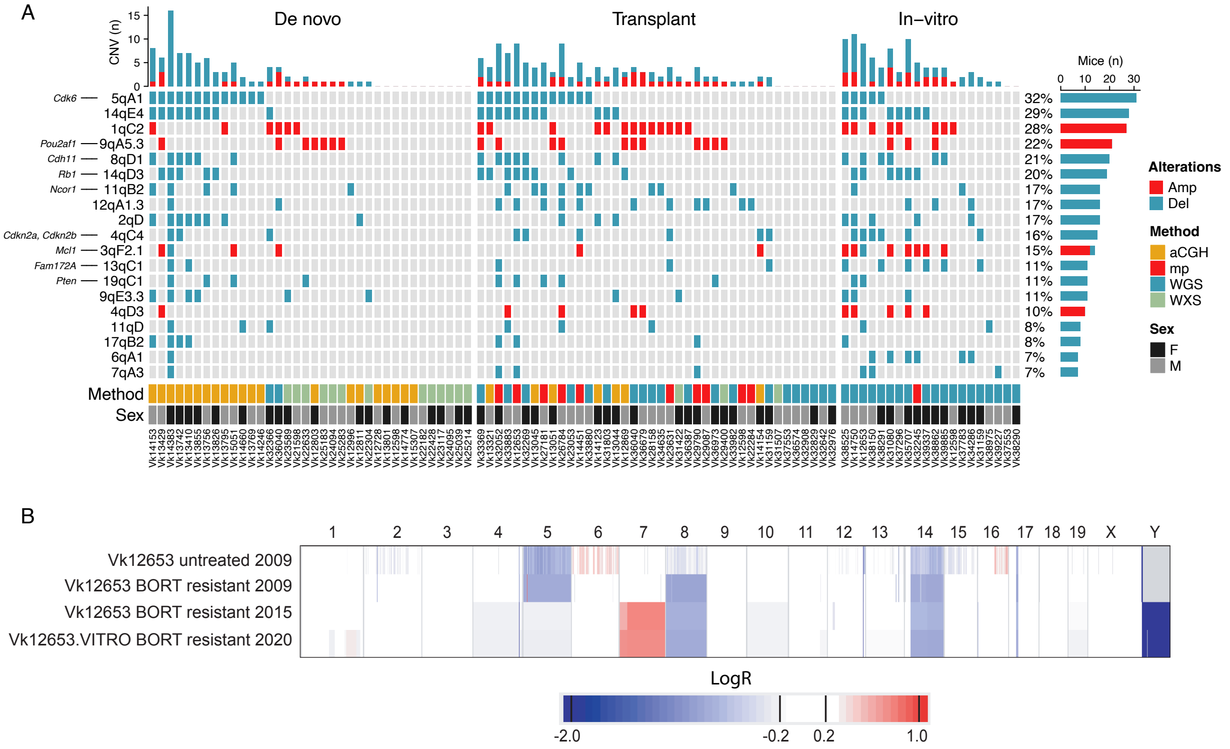

**Supplementary Figure 4. A)** Graphic representation of the Vk*MYC construct (not to scale), with head to tail integration. Green squares represent the kappa variable region and human MYC exons. The position of the two LoxP sites flanking the 3' Kappa enhancer is shown. Horizontal arrow indicates the transcription start point. Arrowhead at the bottom show the position and orientation of PCR primers. **B)** Detection of floxed versus unfloxed Vk*MYC allele by competitive qPCR using primers 1727, 1728 and 1875, shown in A), performed on spleen and BM harvested from Vk22284 tumor bearing mice at the indicated time before and after tamoxifen treatment. **C)** The relative abundance of floxed versus unfloxed Vk*MYC allele after tamoxifen treatment, as shown in B), determined by qPCR using the Δct method.

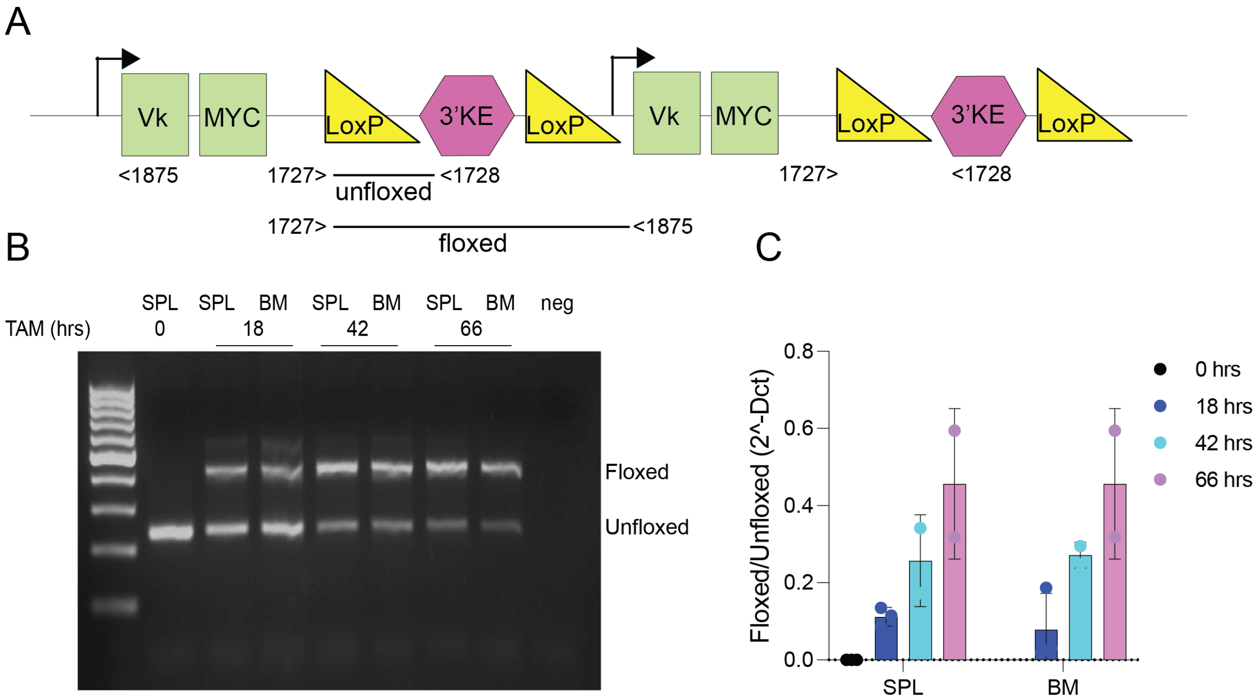

**Supplementary Figure 5. AID and somatic hypermutation activity in Vk*MYC and human multiple myeloma.** SBS= single base substitutions.

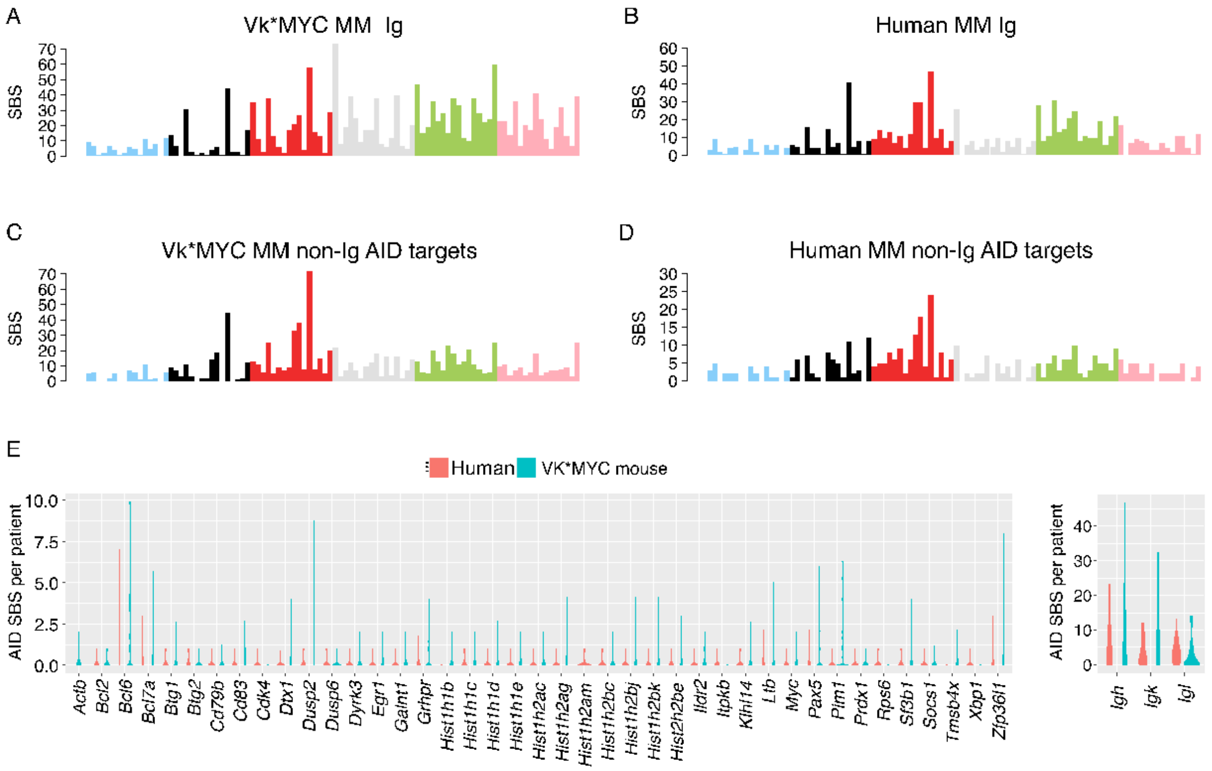

**Supplementary Figure 6. APOBEC mutational contribution and expression in whole exome sequencing (A) and RNAseq data (B) from Vk*MYC multiple myeloma.**

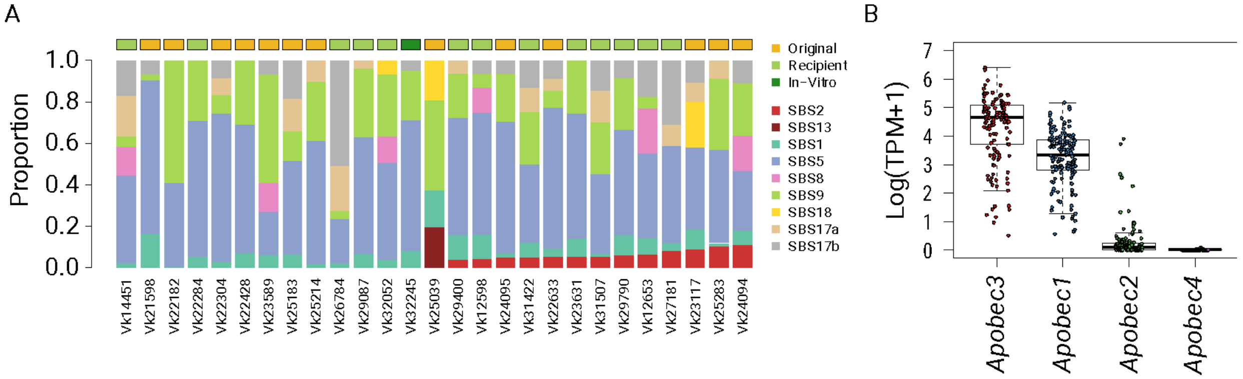

**Supplementary Figure 7. scRNA of 15 Vk*MYC multiple myeloma.**

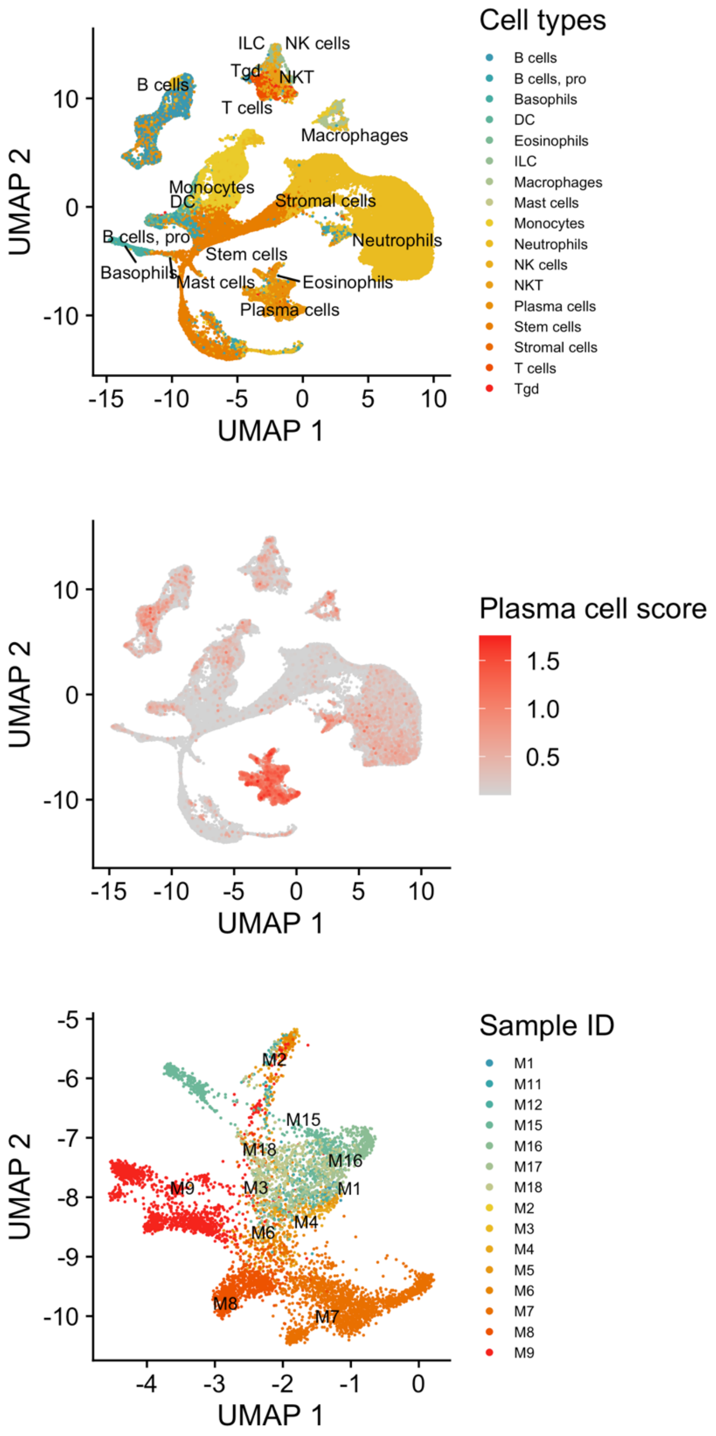
